# Treatment with an Antigen-Specific Dual Microparticle System Reverses Advanced Multiple Sclerosis in Mice

**DOI:** 10.1101/2022.03.25.485841

**Authors:** Alexander J Kwiatkowski, Eric Y Helm, Joshua M Stewart, Theodore T Drashansky, Jonathan J Cho, Dorina Avram, Benjamin G Keselowsky

## Abstract

Antigen-specific therapies hold promise for treating autoimmune diseases such as multiple sclerosis while avoiding the deleterious side effects of systemic immune suppression. In this study, an antigen-specific dual-sized microparticle (dMP) immunotherapy reversed hind limb paralysis when administered in mice with advanced experimental autoimmune encephalomyelitis (EAE). Treatment reduced central nervous system (CNS) immune cell infiltration, demyelination and inflammatory cytokine levels. Mechanistic insights using single-cell RNA sequencing showed that treatment impacted the MHC II antigen presentation pathway in dendritic cells, macrophages, B cells and microglia, not only in the draining lymph nodes, but strikingly also in the spinal cord. CD74 and cathepsin S were among the common genes downregulated in most antigen presenting cell (APC) clusters, with B cells also having numerous MHC II genes reduced. Efficacy of the treatment diminished when B cells were absent, suggesting their impact in this therapy, in concert with other immune populations. Activation and inflammation were reduced in both APCs and T cells. This promising antigen-specific therapeutic approach advantageously engaged essential components of both innate and adaptive autoimmune responses, and capably reversed paralysis in advanced EAE without the use of broad immunosuppressant.

**Significance Statement:** Multiple sclerosis (MS) is a debilitating autoimmune disease that can lead to paralysis. We demonstrate an antigen-specific microparticle treatment can reverse hind limb paralysis when administered in advanced EAE. Single-cell RNA-sequencing and flow cytometry analysis provide evidence the treatment acts by diminishing Ag presentation in APCs, including B cells in the CNS and the draining lymph nodes. Thus, the antigen-specific dual-sized microparticle treatment is a promising therapy even in advanced EAE, and potentially MS.

## Introduction

Multiple sclerosis (MS) is an autoimmune disease where infiltrating immune cells destroy the myelin sheath surrounding axons, leading to defunct neuronal signaling (1, 2). Pro-inflammatory T helper type (Th) 1 and Th17 cells are two of the critical drivers of disease pathology via production of neurotoxic cytokines. These cytokines include granulocyte-macrophage colony-stimulating factor (GM-CSF) and interleukin (IL)-17α (3–5). This pathology is consistent in both human disease and a mouse model of MS, known as experimental autoimmune encephalomyelitis (EAE) (6).

MS does not have a cure, and treatments are largely ineffective or involve systemic immune suppression. Broad-spectrum immunosuppressants are not viable for long-term disease management due to off-target effects, inconsistent disease control and vulnerability to opportunistic infections (7–9). Other treatments such as glatiramer acetate and interferon (IFN)-β are not broadly immunosuppressive but do not slow disease progression (10, 11). B cell depletion with anti-CD20 therapy (Ocrelizumab) is the only current treatment that slows disease progression (12), despite MS pathology being generally believed to be T cell mediated. Thus, MS has a complex pathology with multiple potential therapeutic targets.

To alleviate concerns related to generalized immune suppression, biomaterials fabricated as nano and microparticles (MPs) can co-encapsulate or co-deliver immunomodulatory factors together with specific antigens (Ag) and provide a peripheral tissue-localized targeted therapeutic. Particle-based approaches for EAE have been reviewed (13), with several formulations delivering auto-Ag to reprogram the immune response in autoimmune diseases (14–16). To this end, we employed a dual-sized MP (dMP) system to treat EAE in an Ag-specific manner using Ag-loaded MPs in combination with MPs encapsulating factors for immune cell recruitment and induction of a suppressive phenotype (17–22). The dMP system utilizes four different polylactic-co-glycolic acid (PLGA) MPs of two different sizes, capitalizing on the size limit of phagocytosis to direct the targeted localization of immunomodulatory factors. The two non-phagocytosable MPs (∼50 µm) encapsulate GM-CSF and transforming growth factor-beta one (TGF-β1), while the two phagocytosable (∼1 µm) MPs encapsulate Vitamin D3 (VD3) and the disease relevant Ag, myelin oligodendrocyte glycoprotein_35-55_ (MOG). The control formulation contains the non-specific peptide Ag ovalbumin_323-339_ (OVA). When used in the dMP formulation and injected into peripheral tissue, GM-CSF acts as a recruitment chemokine and dendritic cell (DC) growth factor, while TGF-β1 and VD3 create a suppressive phenotype in recruited cells (23–26). Our previous work showed that subcutaneous injection of dMP treatment before disease onset blocks EAE progression, partly due to a reduction in pathogenic T cells in the central nervous system (CNS) (17).

The present study employs the dMP MOG therapy administered in advanced EAE, with complete hind limb paralysis. Treatment with dMP MOG reversed paralysis, but the non-specific Ag dMP OVA failed to have therapeutic efficacy. The dMP MOG treatment resulted in reduced demyelination and cellular infiltration in the CNS and lowered levels of pro-inflammatory cytokines, including GM-CSF, IL-1β, IL-6, Tnf-α and IL-17α. Mechanistic transcriptomic analysis by single-cell RNA sequencing (scRNAseq) revealed a decrease in MHC II processing and presentation by APCs, including by B cells, both in the draining LNs and in the spinal cord, and additionally by microglia in the spinal cord.

## Results

### Ag-specific dMP MOG formulation administered in mice with advanced EAE reversed hind limb paralysis

Given the success of the Ag-specific dMP MOG treatment before disease onset (17), we administered subcutaneously two doses of the dMP treatment in advanced EAE: the first when mice reached complete hind limb paralysis (score of 3), and the second three days later (Fig. 1A). Ag-specific dMP MOG formulation, but not dMP OVA treatment, reversed disease progression to an average scores of 1 (limp tail) for over 10 days post treatment initiation and overall diminished cumulative scores (Fig. 1B). Thus, dMP MOG treatment, but not a non-specific Ag formulation, is highly effective at reversing hind limb paralysis when administered in advanced EAE.

**Figure 1.**
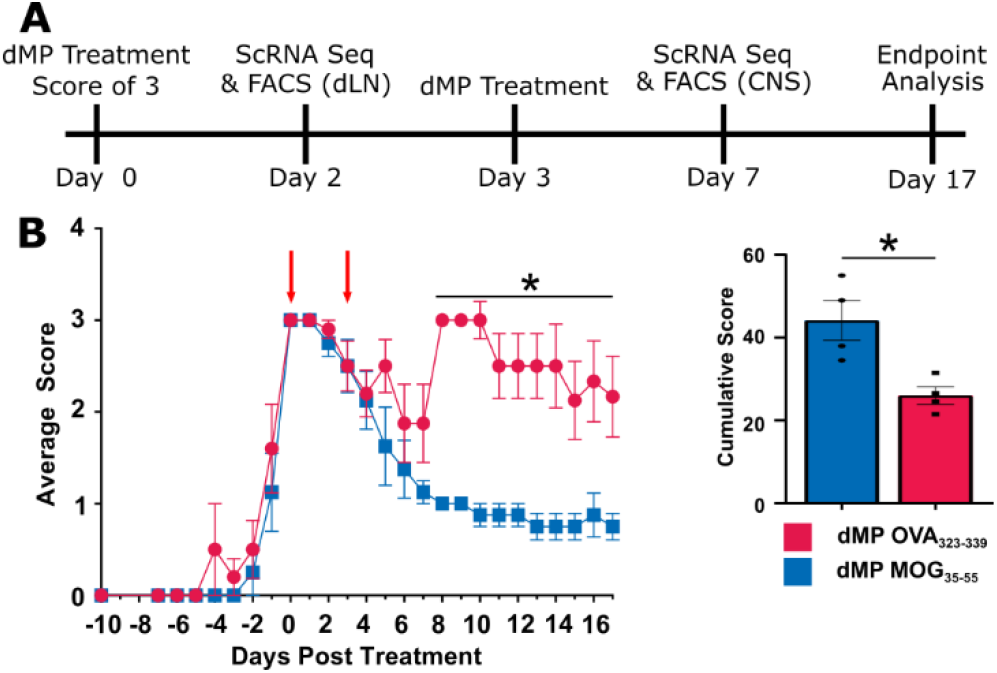
dMP MOG subcutaneous treatment in advanced EAE reverses hind limb paralysis and shows lasting efficacy. a) Timeline of treatments and analysis. b) EAE score curve and cumulative scores following subcutaneous dMP injection initiated at a score of 3 (complete hind limb paralysis) with a second injection three days later. Clinical scores from 0-4: 1: limp tail, 2: weak hind limbs, 3: hind limb paralysis, 4: complete hind limb paralysis and partial hind limb paralysis. Scoring took place every other day until the first mice showed scores, at which point scoring took place daily. The cumulative score took the sum of the scores following treatment and represents the disease severity over the course of the experiment. Score curves were representative of three experiments. n=4-5/group in each experiment * represents p < 0.05

### dMP MOG treatment reduced immune cell infiltration, demyelination and inflammatory cytokines in mice with advanced EAE

Histopathology and flow cytometry analyses of the lumbar spinal cord on day 17 post-treatment showed reduced immune cell infiltration in mice treated with dMP MOG compared to dMP OVA (Fig. 2A-B). In addition, demyelination and large lesions were concomitantly decreased on both days 7 and 17 following dMP MOG treatment compared to dMP OVA (Fig. 2 C-D).

**Figure 2.**
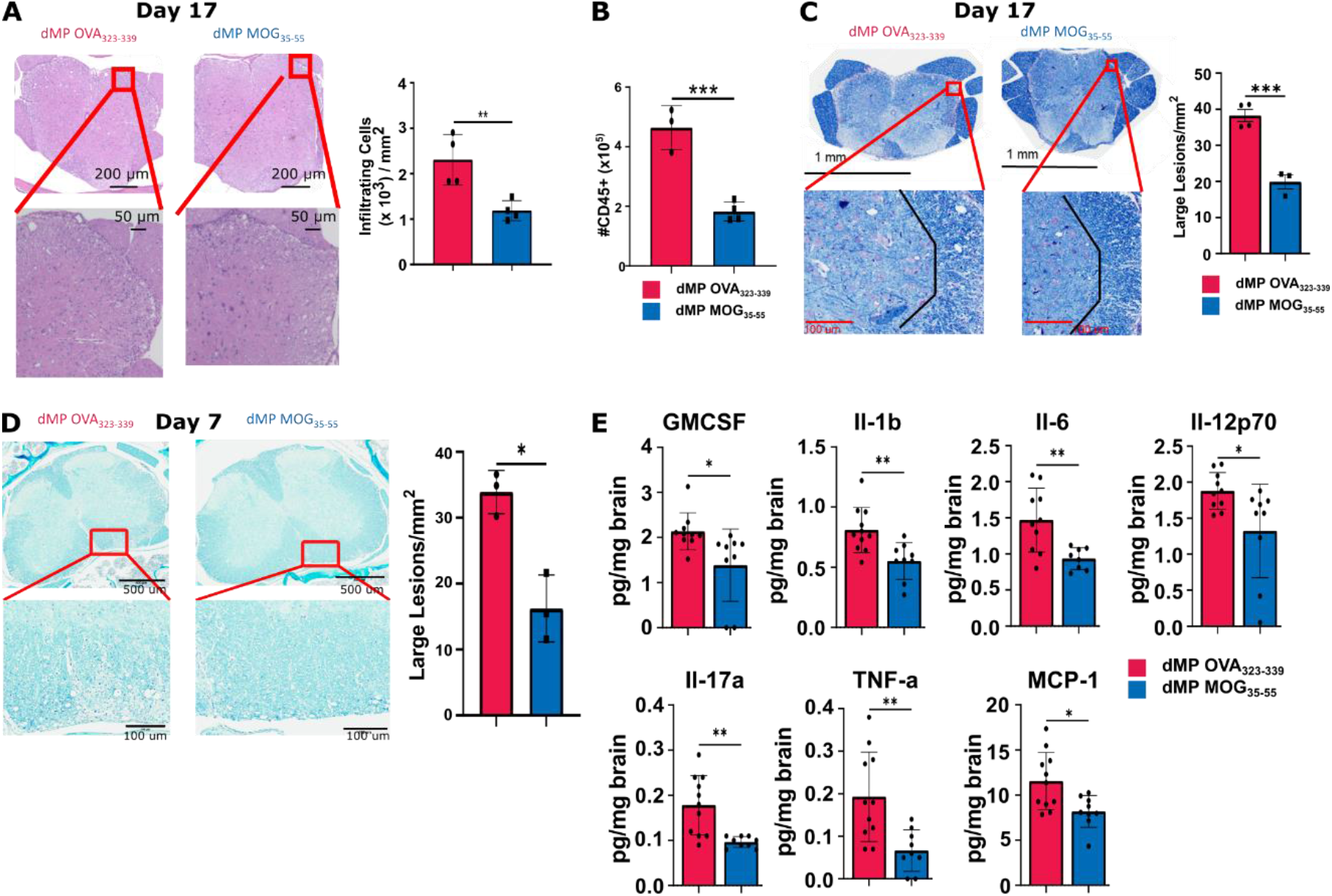
dMP MOG treatment in advanced EAE reduced demyelination, immune infiltration and inflammatory cytokine levels in the CNS. Mice were induced with EAE and treated at a score of 3, as indicated in Fig. 1A, with a second injection three days later. a) Representative images of H&E stained sections and quantification of infiltrating immune cells in the lumbar spinal cord on day 17 post-treatment. Cells were counted and normalized to the area to calculate cells/mm^2^. n=4/group. b) Quantification of CD45+ CNS infiltrates by flow cytometry on day 17 post-treatment. N=3-4/group c) Representative images and quantification of demyelination using luxol fast blue staining in the lumbar spinal cord on day 17 post-treatment. Lesions were counted and those with an area > 400 µm^2^ were classified as a large. n=3-4/group d) Representative images and quantification of demyelination using luxol fast blue staining in the lumbar spinal cord on day 7 post-treatment. n=3 e) Luminex xMAP technology for multiplexed quantification of Mouse cytokines, chemokines, and growth factors from complete brain homogenate at a concentration of 4 mg/mL on day 7 post treatment. n=8-11. d= days post treatment * represents p < 0.05, ** represents p < 0.01

Further evaluation of CNS inflammatory cytokines, known to be associated with MS or pathogenic Th17 cells (1, 3, 4, 27) revealed reduced levels of GM-CSF, IL-1β, IL-6, IL-12p70, IL17α, TNF-α and MCP-1 (CCL2) in dMP MOG treated mice (Fig. 2E). Notably, GM-CSF, IL-17α and TNF-α are produced by Th17 and Th1 cells, the primary pathogenic T cells in MS (28, 29), while IL-6 and IL-1β promote Th17 cell differentiation (30). Increased MCP-1 is known to promote trafficking to the CNS (31). Other cytokines, including IFN-γ or anti-inflammatory IL-10, did not show differences (Fig. S1). Thus, the dMP MOG formulation administered in advanced EAE greatly reduced the hallmarks of MS pathology, namely demyelination, immune cell infiltration and inflammatory cytokine levels in the CNS.

### Common and distinct immune cell clusters were identified in the dLNs and spinal cord of mice with advanced EAE treated with dMP MOG versus dMP OVA

To further determine the mechanisms of action of the dMP MOG treatment, scRNAseq was conducted in the draining inguinal lymph nodes (dLNs) and the spinal cord on day 2 and day 7, respectively, post treatment initiation (Fig. 1A). Sequencing was conducted on total CD45.2 cells from the spinal cord. However, given the reduced frequencies of CD11b+ CD11c+ and CD11b+ CD11c- populations in the dLNs, in order to improve the sensitivity for these two groups, immune cells were sorted into four populations: (1) TCR-β+, (2) CD19+, (3) CD11b+ CD11c+ and (4) CD11b+ CD11c-, and further recombined to 1:1:2 ratio with approximately equal proportions of population 3 and 4.

Cluster definition using Seurat analysis (32) and transcriptomic signatures of hallmark genes identified 18 distinct immune cell clusters in the dLNs and 16 in the spinal cord (Fig. 3A, D). Many clusters overlapped between the dLNs and spinal cord, with tissue specific clusters as well. APCs in the dLNs included one macrophage and three DC clusters (pDCs, cDC1s, cDC2s), while the spinal cord had pDCs, microglia and MHC II^High^ cells (Fig. 3B-C, E-F). MHC II^High^ cells comprised both Itgax^hi^ Adgre1^lo or −^ (DCs) and *Adgre1*^hi^ cells, primarily macrophages (Fig. 3E-F). *Tmem119* and *P2ry12* defined the microglial cluster (Fig. 3E-F). Additionally, both the dLNs and spinal cord had Ly6c^Hi^ and Ly6c^Lo^ monocytic clusters (Fig. 3B-C, E-F). However, dLNs comprised only inflammatory and suppressive neutrophils, identified by the hallmark genes *IL-1β* and *Cd177*, respectively, while the spinal cord contained an additional cluster, namely an immature neutrophil cluster (Fig. 3B-C, E-F).

**Figure 3.**
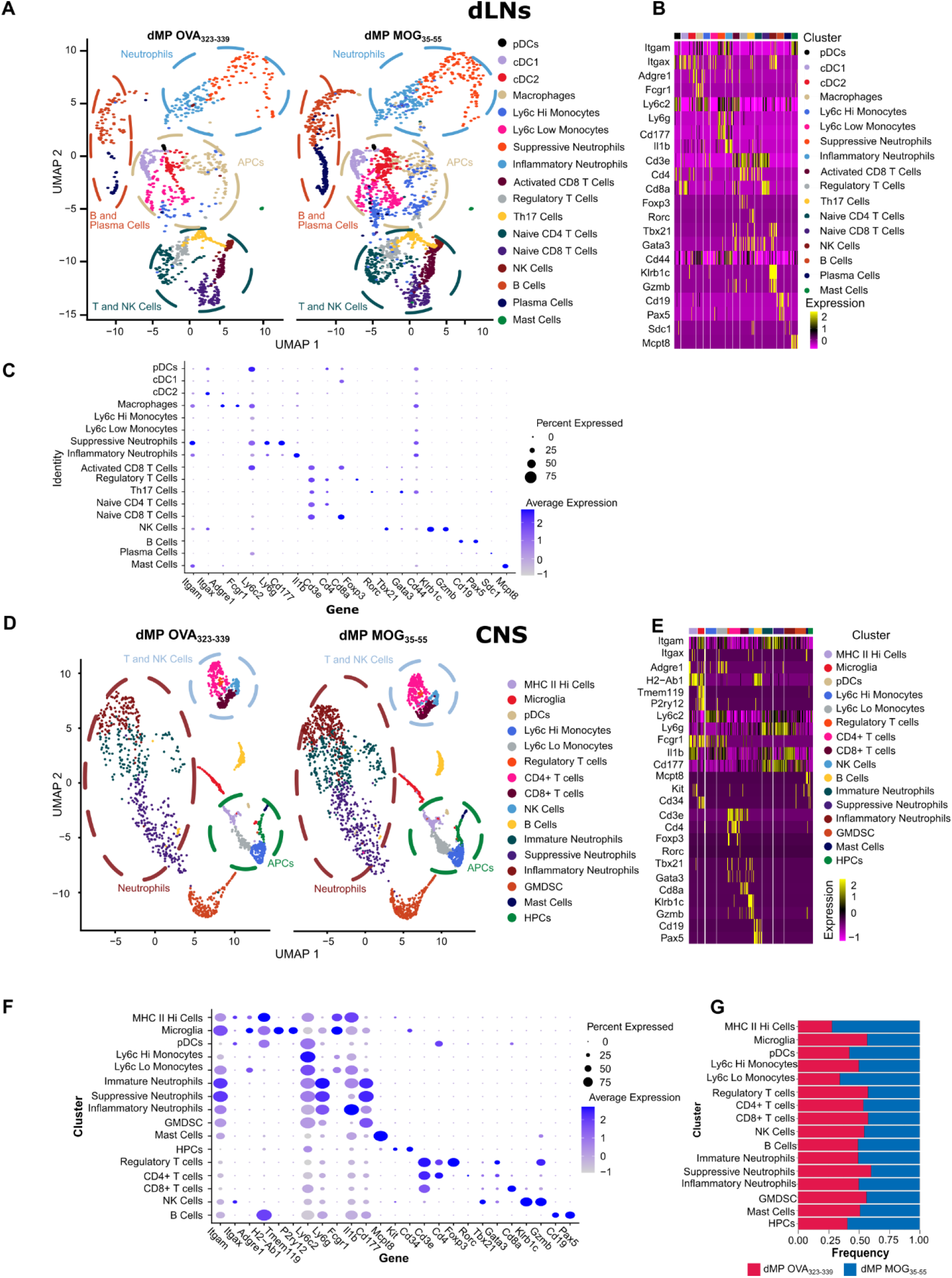
dLNs and CNS scRNAseq cluster identification in mice with advanced EAE treated with dMP MOG versus dMP OVA subcutaneously. a) Side by side uMAP plot showing clusters in dLNs of dMP OVA and dMP MOG spinal cord on day 2. Each dot represents an individual cell where approximately 10,000 cells were analyzed per group with plots downsampled to present more distinct clusters. Circles highlight the location of distinct cell types including cDC1s, cDC2s, neutrophils, T cells and NK cells. Clustering utilized Seurat for initial clustering then hallmark genes to remove non-biologically relevant clusters. b, c) Hallmark genes used to identify each cluster. d) Side by side uMAP plot showing clusters in dMP OVA and dMP MOG spinal cord. Cells were collected from one mouse in each treatment group on day 7 at a score of 3 (Fig. 1A). Each dot represents an individual cell where approximately 10,000 cells were analyzed per group with plots downsampled to clusters more distinctly. Circles highlight the location of distinct cell types including, MHC II^high^ cells, microglia, neutrophils and T cells, NK cells. Clustering utilized Seurat for initial clustering then hallmark genes to remove over clustering. e, f) Hallmark genes used to identify each cluster. g) Fractions of each cluster following post sort recombination as a comparison between dMP OVA and dMP MOG_35-55_. Genes are grouped to form a diagonal of cell type and hallmark genes on the dot plot.

Within the T cell clusters, dLNs contained both activated (*Cd44*+) and naïve CD8+ T cells (Fig. 3A-D), while in the spinal cord, all CD8+ T cells were *Cd44+* (Fig. 3E-F). The dLN CD4+ T cell clusters comprised naïve CD4+, Th17 and T regulatory (Treg) cells (Fig. 3B-C), while in the spinal cord, only CD4+ and Treg cells were identified (Fig. 3E-F). The spinal cord CD4+ T cell cluster contained cells that expressed either *Rorc*, *Tbx21* or *Gata3* (Fig. 3E-F).

*Cd19* and *Pax5* expression defined B cells (Fig. 3B-C, E-F), while plasma cells, classified by expression of *Cd138* (syndecan1), were present only in the dLNs, but not in the spinal cord (Fig. 3B-C). In addition to these populations, *Cd34* expression identified a hematopoietic precursor population present in the spinal cord, but not the dLNs (Fig. 3E-F).

### Genes associated with Ag-presentation are downregulated both in the dLNs and spinal cord APCs of mice with advanced EAE treated with dMP MOG_35-55_

Differential expression analysis of the spinal cord and dLN APC clusters, including B cells and microglia (in the spinal cord), revealed common downregulation of the MHC II Ag presentation pathway (Fig. S2 A, C-E, G-H). *Cd74*, a critical MHC II chaperone protein (33) and cathepsin S (*Ctss*), an enzyme involved in the processing of the MHC II invariant chain (34), were downregulated in numerous dLN and spinal cord APCs, including B cells and microglia (CNS) (Fig. 4A-H). Flow cytometry analysis confirmed a significant CD74 downregulation in the spinal cord DCs (CD11b+CD11c+), macrophages (F4/80+) and microglial (Tmem119+) (Fig. 5A-C). Numerous MHC II genes were downregulated in B cell clusters from both spinal cord and dLNs, in addition to CD74 and Ctss (Fig. 4D, H). Flow cytometry analysis analysis confirmed the reduction in B cell MHC II^high^ population on day 7 post-treatment (Fig. 5D). Multiplex immunofluorescent staining further confirmed a significant reduction in immune cell infiltration (Fig. 5E). In addition, fewer T cells colocalized with APCs, defined as CD74+ (Fig. 5E). In contrast to dMP OVA treated mice, the microglia of dMP MOG treated mice did not show retraction of processes or expression of CD74 (Fig. 5E), suggesting a less active state (35, 36). Furthermore, in agreement with reduced lesion burden (Fig. 2 C-D), astrocytes (GFAP+), known to be numerically elevated in association with larger lesions (37), had reduced numbers in the spinal cord of dMP MOG treated mice compared to dMP OVA control mice (Fig. 5E). Thus, these results altogether demonstrate that the dMP MOG treatment in mice with advanced EAE results in reduced MHC II Ag presentation both in dLN and spinal cord APCs, including microglia and B cells.

**Figure 4.**
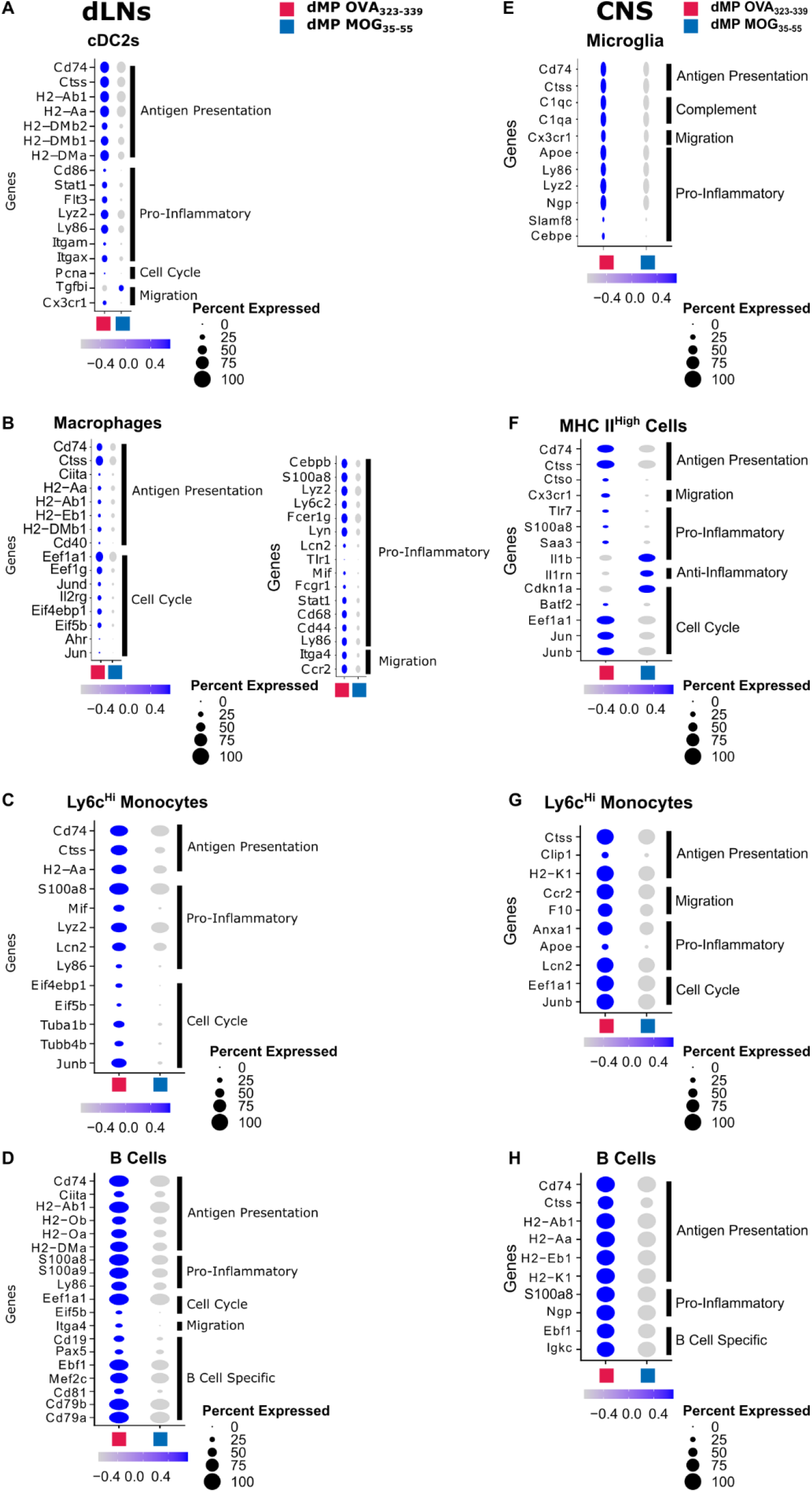
Antigen presenting cells in the dLNs and CNS show a reduction in MHCII Ag presentation and inflammatory gene signature following dMP MOG treatment in mice with advanced EAE. Dot plots highlighting antigen presentation, pro-inflammatory, cell cycle and migration genes differentially expressed in a) cDC2s, b) macrophages, c) Ly6c^Hi^ monocytes and d) B cells in the dLNs on day 7. Dot plots highlighting antigen presentation, pro-inflammatory, cell cycle and migration genes differentially expressed in e) microglia, f) MHC II^High^ cells, g) Ly6c^Hi^ Monocytes and h) B cells in the spinal cord on day 7. Of note are Ag presentation, pro-inflammatory, anti-inflammatory, cell cycle and migration genes. Genes with a p_adjusted_ <0.1 of were considered significant. n=2 mice per group in the dLNs and n=1 mouse per group in the CNS.

**Figure 5.**
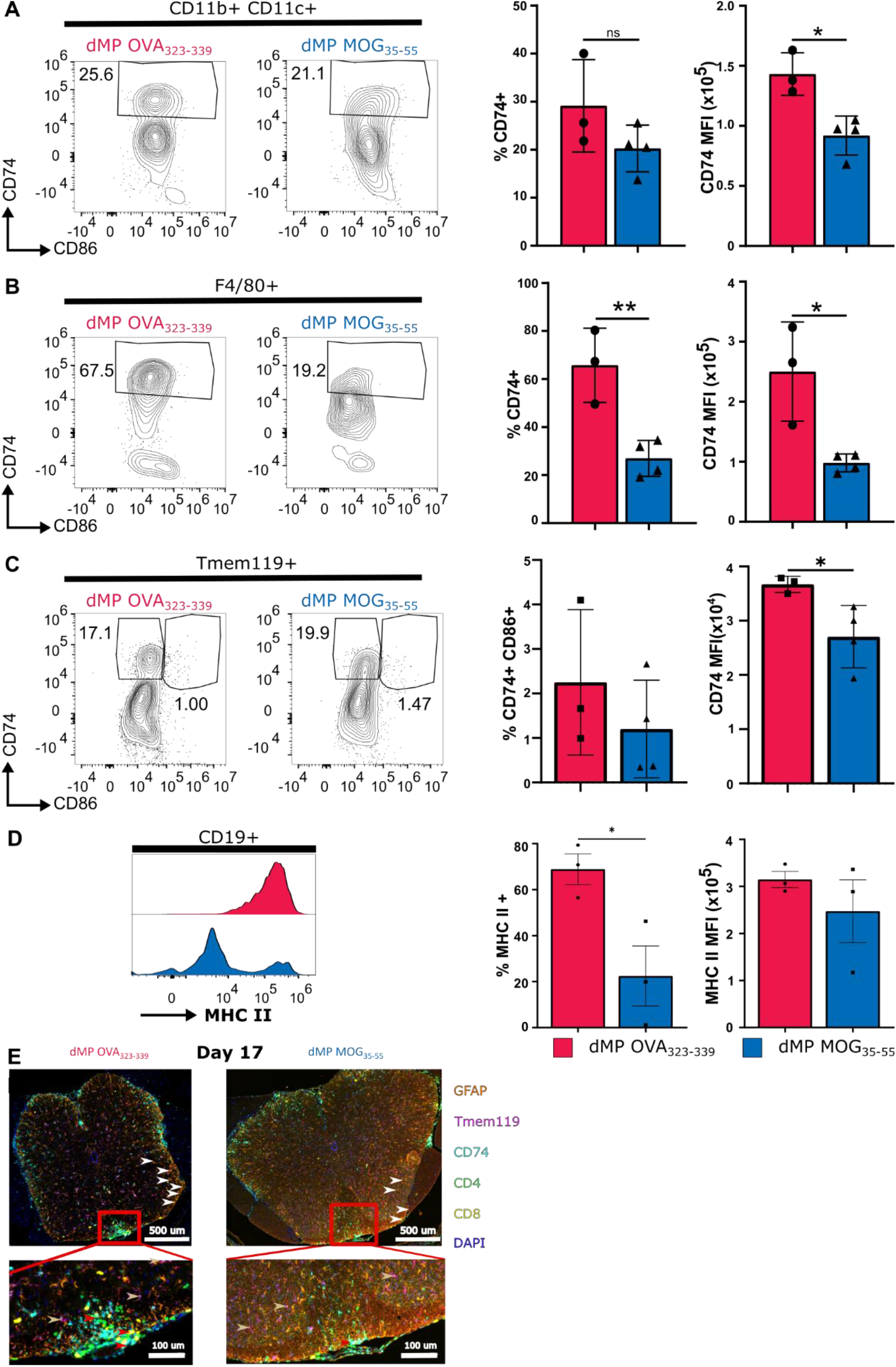
dMP MOG treatment in mice with advanced EAE results in reduction in MHCII components in microglia and CNS DCs, macrophages and B cells. Representative flow plots and quantification of CD74 in the spinal cord on day 7 in a) DCs; b) macrophages; c) microglia. d) MHC II in by B cells on Day 7. e) Multiplex immunofluorescence image of the lumbar spinal cord on day 17. Markers are as follows: CD4 (T helper cells) in green, CD8 (cytotoxic T cells) in yellow, CD74 (MHC II chaperone protein) in cyan, Ki67 (proliferation marker) in red, GFAP (astrocytes) in orange and Tmem119 (microglia) in magenta. Red arrows show APCs interacting with T Cells, white arrows point to astrocytes and gold arrows point to microglia. The dorsal column in the red box is magnified in the bottom image. All populations were pregated on live CD45+ DCs and Macrophages. B cells were negative for Tmem119 and CD3e. Macrophages and DCs were negative for B220/CD19, and macrophages were negative for CD11c. n=3-4/group * represents p<0.05 ** represents p <0.01

### B cells are partially responsible for dMP MOG treatment efficacy

Given the numerous changes in the B cell clusters and the recent success with anti-CD20 therapy in MS, we further investigated the impact of B cells on dMP MOG treatment efficacy in Mumt^−/−^ mice, which are deficient in B cells (38, 39). Previous publications showed that B cell absence had no significant impact on EAE scores (39). Our results show that normalized cumulative scores following the dMP OVA treatment were comparable between C57BL/6 and MuMt^−/−^ mice (Fig. 6A). Thus, consistent with the previous observation that the absence of B cells does not impact EAE in C57BL/6 induced with MOG_35-55_ (39). Mumt^−/−^ mice induced with EAE and treated with dMP MOG had lower scores than dMP OVA and overall lower cumulative scores (Fig. 6B). However, when comparing normalized cumulative scores, C57BL/6 mice treated with dMP MOG outperformed Mumt^−/−^ mice treated with dMP MOG (Fig. 6C), suggesting that B cells are an essential component of the dMP MOG efficacy, with the B cell Ag-presentation pathway likely implicated (Fig. 4D, H and 5D). Given that the scores were still reduced, comparing dMP MOG versus dMP OVA treated Mumt^−/−^ mice, this suggests that other immune populations also play a major role in addition to B cells.

**Figure 6.**
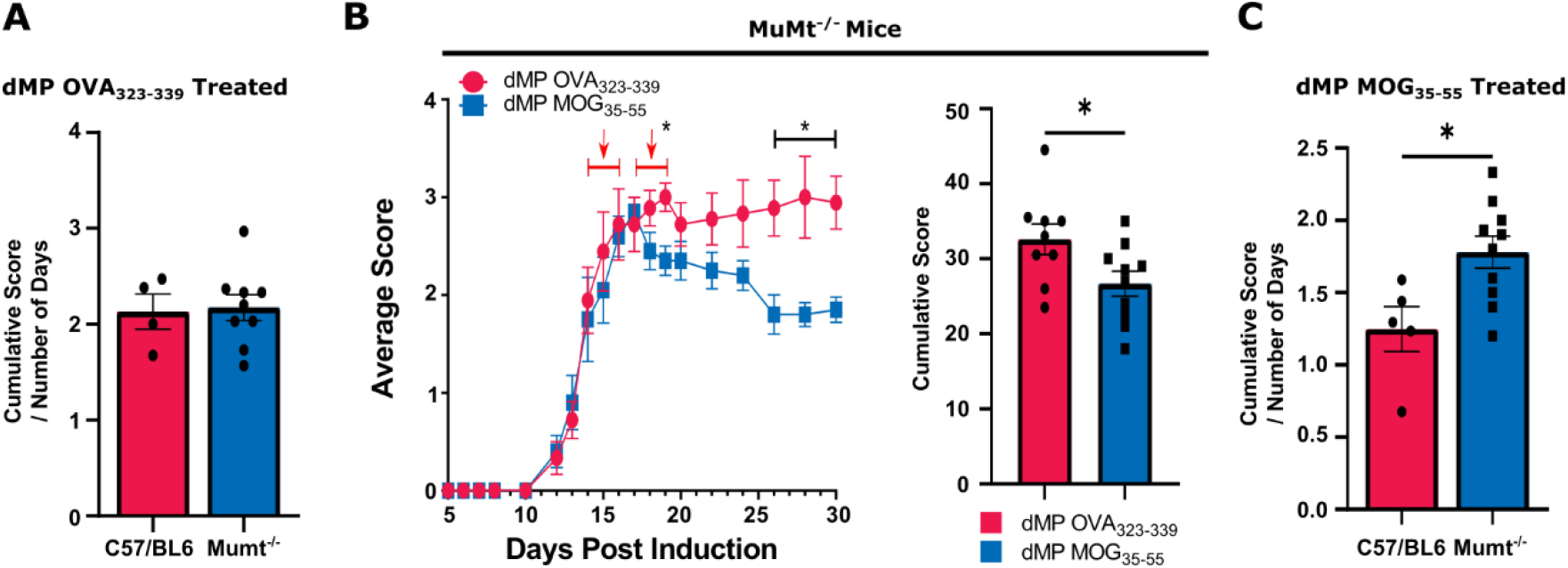
B Cells are partially responsible for dMP MOG treatment efficacy. a) Comparison of cumulative score normalized by the number of days post treatment initiation in EAE C57BL6 and EAE muMt^−/−^ mice administered with dMP OVA. b) EAE score curve and cumulative scores following treatment initiated at a score of 3 in muMt^−/−^ mice as in Fig. 1A, with a second injection three days later. c) Comparison of cumulative score normalized by the number of days post treatment initiation in C57BL6 and muMt^−/−^ mice administered with dMP MOG. n= 5-9 per group and represents 2 independent experiments* Represents p< 0.05

### Genes associated with inflammation are downregulated in APCs and T cells of EAE mice treated with dMP MOG

Both APCs and T cells in the spinal cord and dLNs showed reduced mRNA belonging to inflammatory pathways following dMP MOG treatment (Fig. S2, S3). Although not all changes were present in all APC clusters, key genes associated with inflammation, activation, maturation and cytokine production (33, 40–42) were downregulated in the dLNs and spinal cord, including *Lyz2, Lcn2, Ly86, S100a8, S100a9, Fcer1g, Mif, Stat1*, *Cebpb, Cd44, Cd68, Cd86, Cd40, Itgam, Itgax, and Flt3* and *Ly6c2* following dMP MOG treatment (Fig. 4A-D). Interestingly, several genes associated with dysfunctional microglia, including *Cebpe, Lyz2*, *Cx3cr1* and *Apoe* (33, 43) or lesion-associated microglia, such as complement genes (44–46), were downregulated in the microglia of dMP MOG treated mice (Fig. 4E). T cells in the dLNs and spinal cord on days 2 and 7 post treatment downregulated a similar gene signature with some distinct genes, including *S100a8, S100a9, Lcn2, Lyz2, Ngp and Anxa1* (Fig. 7A-G).

**Figure 7.**
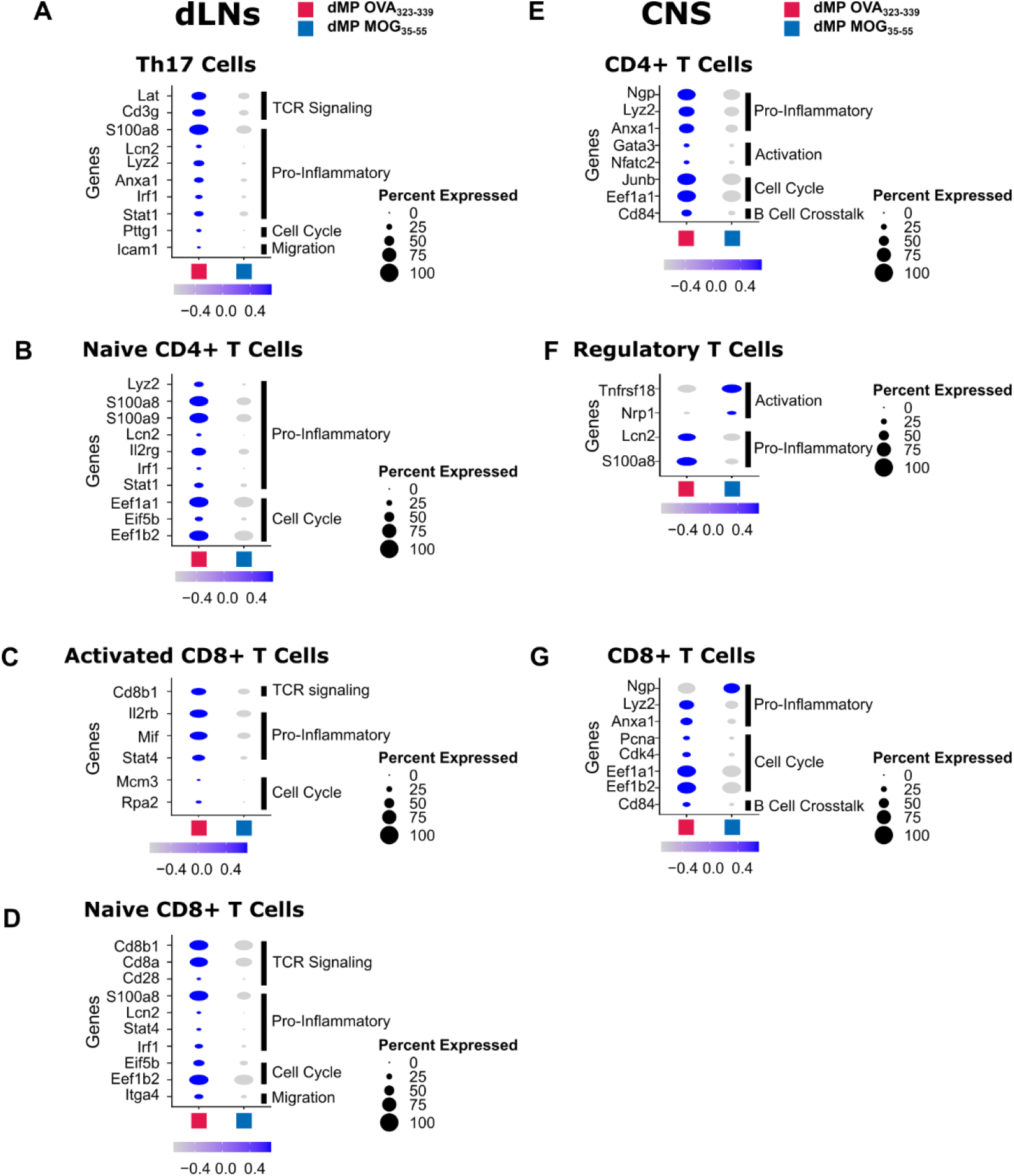
T Cells in dLNs and CNS show a reduction in inflammatory genes following dMP MOG treatment in mice with advanced EAE. b) Dot plots highlighting genes of interest differentially expressed in a) Th17 cells, b) Naïve CD4+ T cells, c) Activated CD8+ T cells and d) Naïve CD8+ T cells in the dLNs on day 2, specifically TCR signaling, pro-inflammatory, cell cycle and migration genes. Dot plots showing pro-inflammatory, activation and cell cycle genes in e) CD4+ T cells, f) Treg cells and g) CD8+ T cells in the spinal cord on day 7. Genes with a p_adjusted_ <0.1 of were considered significant. n=2 mice per group in the dLNs and n=1 mouse per group in the CNS.

In addition to decreased expression of pro-inflammatory genes, Th17 cells and CD8+ T cell clusters in the dLNs showed downregulation of TCR signaling genes (Fig. 7A C-D). In conjunction with reduced Ag presentation by APCs, this suggests a concomitant mechanism contributing to treatment efficacy (Fig. 4A-H, Fig. 7A, C-D, S3A, C). Additionally, Treg cells in the spinal cord also downregulated the inflammatory genes *S100a8 and Lcn2* and showed increased expression of *Tnfsf18* (encodes for Gitr) and *Nrp1* (Fig. 7F), known to be associated with Treg cell activation (47–51). Altogether, results show that dMP MOG treatment reduced pro-inflammatory signature genes in numerous APCs and T cell clusters in the dLNs and spinal cord, which corresponded with reduced production of pro-inflammatory cytokines in the CNS.

### Expression of genes involved cell cycle, DNA replication and translation was downregulated in diverse immune populations of EAE mice treated with dMP MOG

Cell cycle, DNA replication and ribosome pathways were downregulated following dMP MOG treatment (Fig. S2A-C, F-G, S3A-B, E-G, S4, S5). Specifically, dMP MOG treated mice downregulated cell cycle genes including *Mcm3, Cdk4* (Fig. 7C, F) along with DNA replication and repair genes such as *Rpa2, Pcna* and *Pttg1* (Fig. 4A, 7A, C, G). Furthermore, translation genes were also downregulated in the dLNs and spinal cord following dMP MOG treatment. These included *Eif5b, Eef1b2, Eef1a1* (Fig.4C-D, F-H, 7B, D-F). Additionally, proliferation related genes *Jun, Junb and Jund* (52–54) were reduced in dLNs and spinal cord (Fig. 4B-C., 4F-G, 7E). The downregulation of these genes and pathways are consistent with a reduction in immune cell activation following dMP MOG treatment.

## Discussion

Our results show that Ag-specific dMP MOG treatment administered subcutaneously in advanced EAE effectively reversed paralysis, an improvement over many other preclinical treatments, which are typically initiated before EAE induction or at the onset of disease (15, 16, 55–62). Interestingly, despite the administration of the same dose of recruitment and suppressive factors, GM-CSF, TGF-β1 and VD3, only the formulation with disease specific MOG Ag has efficacy, showing the ability of the treatment to induce Ag-specific immune suppression. Mechanistically, the dMP subcutaneous injection depot has been shown to recruit and favor DC differentiation with a suppressive phenotype (17, 19). Our results show that the treatment acts more broadly on APCs. Following trafficking of APCs to the dLNs, dMP MOG treatment resulted in reduction of MHC II Ag presentation, conditioning toward an Ag-specific suppression of T cells with downregulation of pro-inflammatory gene signature in dLN T cells. These changes in T cells implicate a potential feedback loop in the dLNs to further attenuate the autoimmune response. Both APCs and T cells have a less pro-inflammatory phenotype in the spinal cord. Notably, less MHC II Ag-presentation in the spinal cord provides a conserved mechanism between the periphery and primary tissue. Furthermore, we report reduced Ag-presentation and activation in resident microglia and astrocytes, respectively. In conjunction with these changes, reduced levels of the neurotoxic cytokines GM-CSF and IL-17α and reduced demyelination suggest a return toward homeostasis in the CNS. A twin study in MS revealed that genes including *Ccr2* and *Ciita* were elevated in the twin with MS, interestingly these two genes are downregulated following dMP MOG treatment (63). In addition to DCs, B cells show a decrease in MHC II Ag presentation and inflammation in the dLNs and the spinal cord. dMP MOG treatment in C57BL/6 mice outperformed Mumt^−/−^ mice demonstrating the important contribution of B cells in the treatment, even though disease progression in this model is not B cell-dependent. Given the fact that the dMP MOG still had an impact versus dMP OVA on EAE scores in Mumt^−/−^ mice demonstrates the importance of other immune populations in mediating the effect of the treatment. Modulation of B cells provides another prong to the mechanism of dMP efficacy that has not been noted in previous work. Overall, we propose a mechanism by which the Ag-specific dMP treatment halts pathogenesis by decreasing the MHC II Ag presentation pathway, immune activation, reduceing immune cell infiltration in the CNS and hallmark inflammatory cytokine signature (Fig. 8). Wholly, these changes indicate an environment less prone to maintaining Ag-specific T cell activation and overall inflammation, including in the CNS. Delivering treatment in advanced disease shows promise for dMP use as a therapeutic and substituting MOG with other MS-specific Ags or combinations could improve translational capabilities. Altogether, this work highlights the promise of an Ag-specific dMP therapy to reverse paralysis through targeted suppression of autoimmune responses.

**Figure 8.**
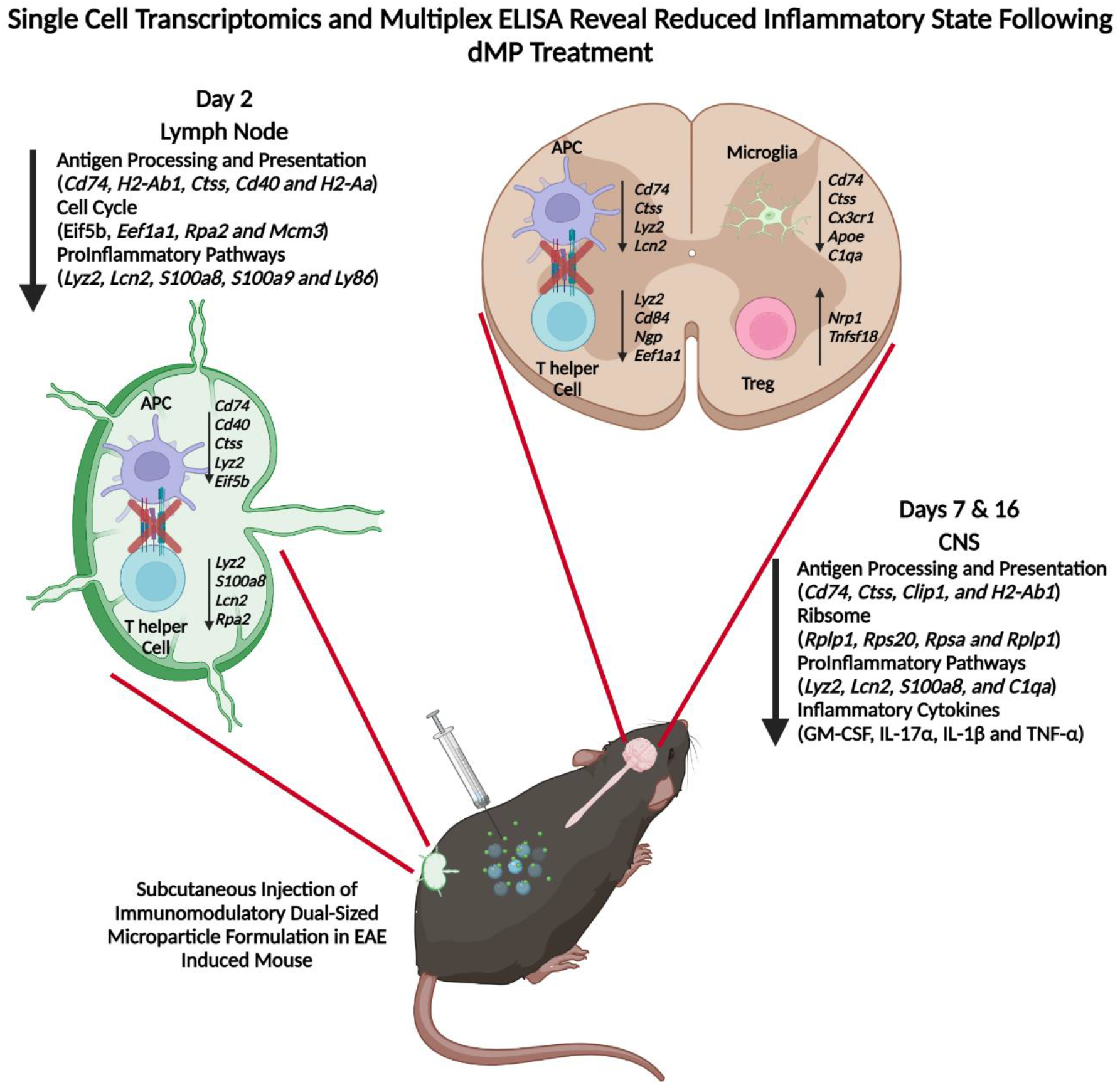
Dual microparticle (dMP MOG) immunotherapy highlighting reprogramming in both the lymph node and CNS.

## Materials and Methods

### Mice, EAE induction, and treatment

Pathogen-free conditions at the University of Florida housed C57BL/6 mice (B6NTac) following purchase from Taconic Biosciences and Mumt^−/−^ mice purchased from Jackson Laboratory. EAE induction utilized 9–13-week-old mice female mice. All experiments were approved by the University of Florida Institutional Animal Care and Use Committee.

EAE induction utilized kits from Hooke Laboratories (Hooke Laboratories Inc., Cat# EK-2110). Subcutaneous injections at both the scruff of the neck and the base of the tail contained 100 μL of MOG_35-55_/Complete Freund’s Adjuvant, resulting in 200 μL injected. 2- and 24-hours following injection of emulsion, 100 μL of pertussis toxin (4 μg/mL) was injected intraperitoneally. Following seven days of incubation, clinical scores were evaluated as follows: 1: flaccid tail, score 2: weak hind limbs, score 3: hind limb paralysis and score 4: quadriplegia.

Treatment consisted of an injection of a cocktail of GM-CSF, TGF-β1, VD3 and Ag-loaded MPs. Subcutaneous MP administration in the center of the back took place at a score of 3, with a second injection three days later.

### Acquiring a single cell suspension from CNS

Following euthanasia, perfusion through the right ventricle with 10 mL of ice-cold PBS cleared circulating cells from the CNS vasculature. A gentleMACS dissociator (Miltenyi Biotec) homogenized the tissue following the brain and spinal cord harvest. Collagenase A (0.33 mg/mL) digestion for 25 minutes shaking at 37 °C followed by mechanical dissociation through a 70 μm filter removed excess CNS tissue. Following centrifugation at 380 G for five minutes, a 30% isotonic Percoll (GE Healthcare), underlaid with 70% isotonic Percoll and spun at 650 G for 25 minutes with no brake separated red blood cells, immune cells and myelin. The interphase between the 30-70% layers contained mononuclear cells, and these were stained for flow cytometry or prepared for single-cell RNA sequencing.

### Staining and flow cytometry

Following Fc blocking for 15 minutes, staining used surface markers (Extended Methods) and fixable viability dye (Affymetrix, Life Technologies) for 30 minutes at 4 °C followed by 2% PFA fixation for 10 minutes at room temperature. Flow cytometry used a BD LSR II with BD FACS DIVA software for data acquisition (BD Biosciences) or Cytek Aurora (Cytek Biosciences) for spectral cytometry. All data analysis used FlowJo (BD Biosciences).

### Histological Analysis

Briefly, spinal cords were harvested with the surrounding vertebrae and placed in 10% neutral buffered formalin overnight. Samples were then decalcified for four hours before processing and embedding. Following processing and embedding, samples were sectioned at 4 µm and stained (Extended Methods).

### Multiplex Cytokine Analysis

Brains were harvested on day 7 post treatment and homogenized using gentlemacs dissociation. Samples were then centrifuged at 400 G for 5 minutes then resuspended in 1% NP 40 Lysis buffer. This solution was placed in a 2 mL blast tube with 500 µL of 800 µm glass beads. A bead blaster lysed the cells and homogenized the tissue. Samples were spun down at 10000G for 10 minutes and a BCA (Thermo Fisher) quantified the protein level. Protein levels were normalized to 4 mg/mL protein before analysis. Eve Technologies used Luminex xMAP technology to conduct cytokine analysis using a Luminex 200 system (Luminex). Eve’s 18-Plex Discovery Assay® (Millipore Sigma) evaluated GM-CSF, IFNγ, IL-1α, IL-1β, IL-2, IL-4, IL-5, IL-6, IL-7, IL-10, IL-12 (p70), IL-13, IL-17A, KC/CXCL1, LIX, MCP-1, MIP-2 and TNF-α according to the manufacturer’s protocol. Sensitivity ranged from 0.06 to 9.06 pg/mL.

### Single-cell RNA-sequencing

scRNAseq used cells from the spinal cords of mice with treatment initiated at a score of 3. After harvesting cells from the spinal cord as previously described, fixed viability dye and CD45.2 staining took place. Following staining, cells sorting collected live CD45.2^+^ cells before processing from scRNAseq. The analysis used CellRanger (10x genomics) in Mobaxterm and Seurat (Satija Lab) in Rstudio conducted cluster analysis and differential expression (Extended Methods).

Fraction calculation involved normalizing the frequency of each cluster to the total number of cells from that mouse followed by comparing the frequencies between groups— a threshold of 65% determined differences in frequency. Hallmark genes include known defining genes including *Cd3e* for T cells, Tmem119 for microglia, *Itgam, Itgax* and *Adgre1* for MHC II^high^ cells and *Ly6c2* and *Ly6g* and *Cd177* and *Il-1b* for neutrophils. Further genes defined the phenotype, including T cell transcription factors *Rorc*, *tbx21*, *Foxp3* and *Cd177* and *Il-1b* for neutrophils.

### Statistical analysis

Mann-Whitney U-Test determined differences in EAE score. Two tailed students t-tests evaluated changes in histological and flow cytometry data. Graphpad Prism 8 was used for analyses.

## Acknowledgment

We would like to acknowledge our funding sources R01 AI33623 to BGK and DA, R21 NS123596 to BGK, R01 AI067846 and R01 AI161845 to DA and T32 DK108736 to AJK. Also grant P30CA076292 to Moffit Cancer Center. The graphical abstract was created with BioRender.com. We would also like to acknowledge Dr. Daryoush Saeed-Vafa, Dr. Carlos Morán-Segura, Neale Lopez-Blanco and Jonathan Nguyen of the Advanced Analytical and Digital Laboratory for assistance with multiplex immunofluorescence staining and imaging. Specifically, Carlos Moran Segura for antibody panel design and multi-spectra scanning, Jonathan Nguyen for image quantification within HALO and Dr. Daryoush Saeed-Vafa for laboratory direction. Also, Noel Clark of the tissue core for assistance with tissue processing and sectioning.

## Supplementary Information

**This PDF file includes:**

Extended Methods

Figures S1 to S5

### Extended Methods

#### Histological Analysis

Using the Leica protocol, samples were dehydrated with ethanol starting with 90% and two rinses of 100% for 15 minutes each. Cleaning then used three xylene washes for 20, 20, and 45 minutes, respectively. Samples were exposed to paraffin wax for three changes of 30, 30, and 45 minutes, respectively. The spinal cord was embedded (Tissue-Tek) so that the L1 vertebrae was the first cross-section cut and ended at L6. The blocks were sectioned at 4 µm and dried overnight before staining. All staining sections were deparaffinized by the following: two five minute changes of xylenes, two three minute changes of 100% ethanol, one minute change of 95% ethanol, and a rinse in deionized water. H&E staining utilized a kit from Vector labs. Briefly, the protocol was 5 minutes in hematoxylin, two rinses of deionized water, bluing reagent for 15 seconds, rinsed in 100% ethanol, 3 minutes in Eosin Y, and three two minute rinses in 100% ethanol then coverslipped. For luxol fast blue staining, after deparaffinization, 0.1% LFB was added to the slides for two hours at 60 °C followed by a three minute change in 70% ethanol and a change in each deionized water, lithium carbonate, and distilled water. Counterstain used cresyl violet for 10 minutes followed by rinses in 95% ethanol and xylenes twice; then the slides were coverslipped. Imaging at 5x and 20x magnification took place on a Zeiss Axio Inverted Microscope (Carl Zeiss).

#### Multiplex Immunofluorescence

Formalin-fixed and paraffin-embedded (FFPE) tissue samples were immunostained using the AKOAYA Biosciences OPAL TM 7-Color Automation IHC kit on the BOND RX autostainer (Leica Biosystems). The OPAL 7-color kit uses tyramide signal amplification (TSA)-conjugated to individual fluorophores to detect various targets within the fluorescent multiplex assay.

Sections were baked at 65 °C for three hours then transferred to the BOND RX (Leica Biosystems). All subsequent steps (ex., deparaffinization, antigen retrieval) were performed using an automated OPAL IHC procedure (AKOYA). OPAL staining of each antigen occurred as follows: heat induced epitope retrieval (HIER) was achieved with EDTA pH 9.0 buffer for 20 minutes at 95°C before the slides were blocked with AKOYA blocking buffer for 10 minutes. Then slides were incubated with primary antibody and one of the OPAL fluorophores during the final TSA step. Staining used the following antibodies, TMEM119 (Abcam, 28-3, HIER- EDTA pH 9.0, 1:100, dye 570), Ki67 (Abcam, SP6, HIER- Citrate pH 6.0, 1:300, dye 620), CD74 (LS Bio, Rb poly, HIER- EDTA pH 9.0, 1:600, dye 520), GFAP (Abcam, EPR1034Y, HIER- EDTA pH 9.0, 1:500, dye 540), CD4 (CST, D7D2Z, HIER- EDTA pH 9.0, 1:100, dye 650), and CD8 (CST, D4W2Z, HIER- EDTA pH 9.0, 1:100, dye 690). All slides were imaged with the Vectra®3 Automated Quantitative Pathology Imaging System. Individual antibody complexes were stripped after each round of antigen detection. After the final stripping step, DAPI counterstain is applied to the multiplexed slide and is removed from BOND RX for coverslipping with ProLong Diamond Antifade Mountant (ThermoFisher Scientific).

#### Antibodies

Cells were stained with the following antibodies: CD11b (BV510, clone: M1/70), CD11c (BV711, clone: N418), F4/80 (PE, PE-Cy7, clone: BM8), MHCII (BV421, APC-ef780, clone: M5/114.15.2), TCR-β (ef450, clone: H57-597), Ly6c (Pacific Blue, clone:HK1.4), Ly6g (PE-Dazzle594, clone: 1A8), CD8α (BV605, SB600, clone: 53-6.7), CD86 (PE-Cy5, APC, clone: GL1), CD4 (BV785, SB780, clone: RM4-5), GR-1 (PE_Cy7, clone: RB6-8C5), CD19 (APC, clone: 1D3), CD45.2 (AF700, clone:104), B220 (BV650, clone: RA3-6B2). Tmem119 (af488, clone: V3RT1GOsz, 28-3), CD74 (AF647, clone: In1/CD74), CD3ε (BV650, clone: 145-2C11), Ki-67 (PE-Dazzle594, clone: 16a8), Foxp3 (PE-cf594, clone: MF23), Rorγt (APC, clone: AFKJS-9), tbet (BV421, clone: 4B10), IL-23R (BV421, clone: 12B2B64), CD16/32 (FCγ III/II receptor, clone 2.4G2), and Fixable viability dye (ef780, Zombie NIR)

**Fig. S1.**
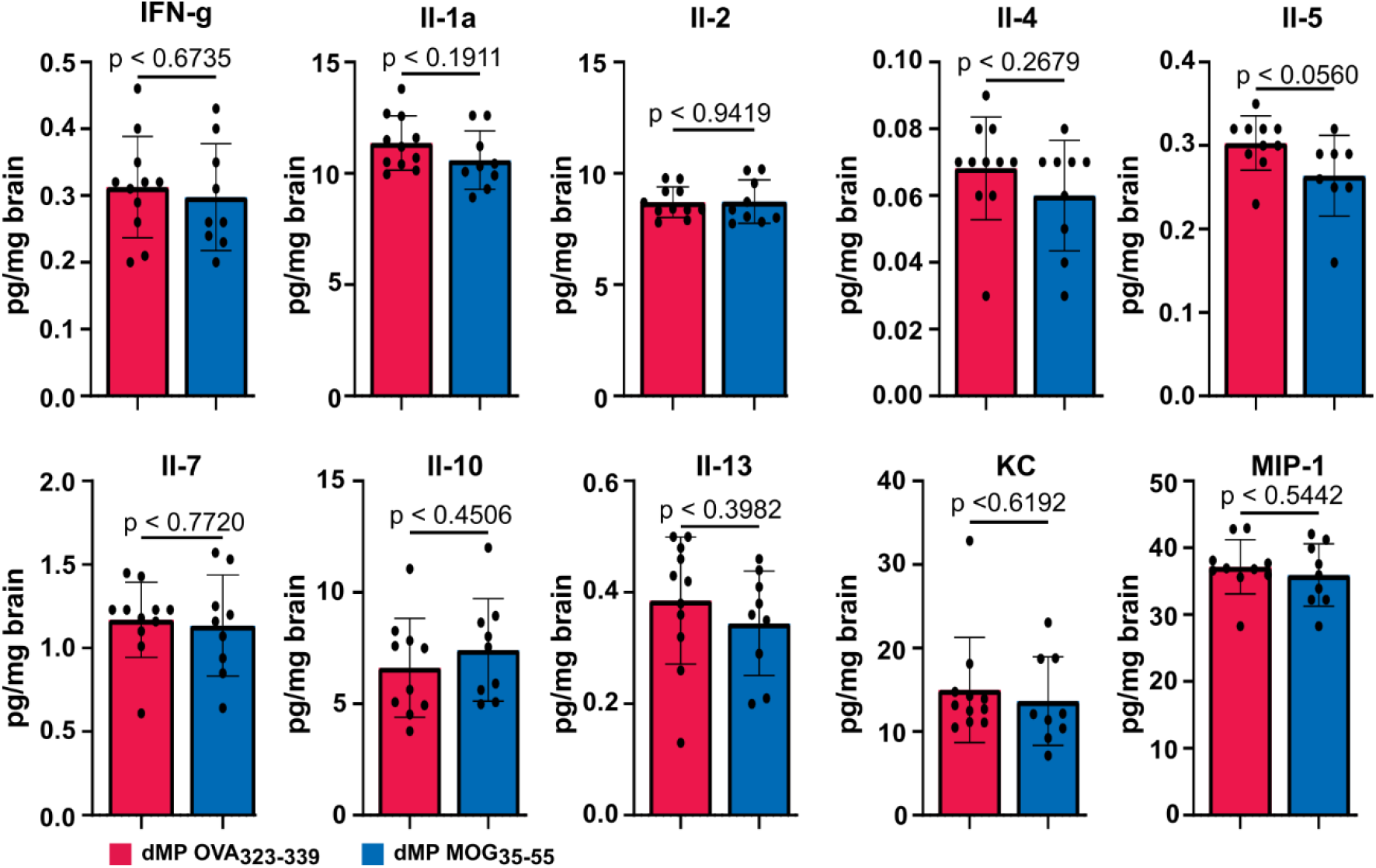
Cytokines less correlated with MS were not differently produced in the CNS between dMP MOG vs. dMP OVA treatment. Luminex xMAP technology multiplexed quantification of Mouse cytokines, chemokines, and growth factors from complete brain homogenate at a concentration of 4 mg/mL on day 7 post treatment. n=8-11 * represents p < 0.05, ** represents p < 0.01

**Fig. S2.**
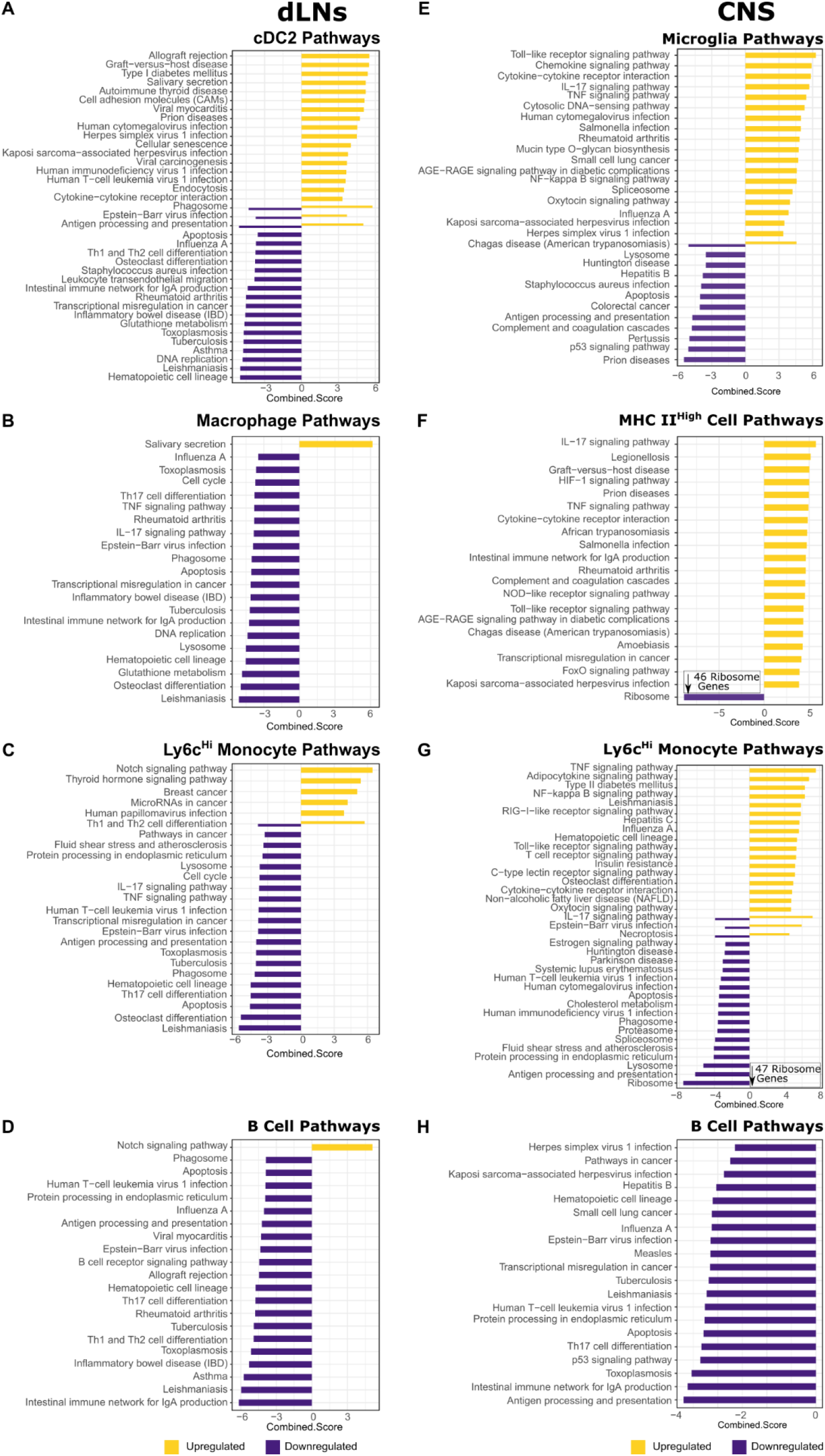
APCs downregulated Antigen presentation, activation and Pro-inflammatory pathway genes in both the dLNs and CNS. Pathways modulated in a) cDC2s, b) macrophages, c) Ly6c^Hi^ monocytes and d) B cells located in the dLNs at day 2, specifically antigen presentation. Pathways modulated in e) microglia, f) MHC II^High^ cells, g) Ly6c^Hi^ monocytes and h) B cells in the spinal cord at day 7, specifically antigen presentation and ribosome. Genes with p_adjusted_ <0.1 were used in pathway analysis. Significant genes were compared to the KEGG pathways and p_adjusted_ <0.1 denoted a pathway was up or downregulated. n=2 mice per group in the dLNs and n=1 mouse per group in the CNS

**Fig. S3.**
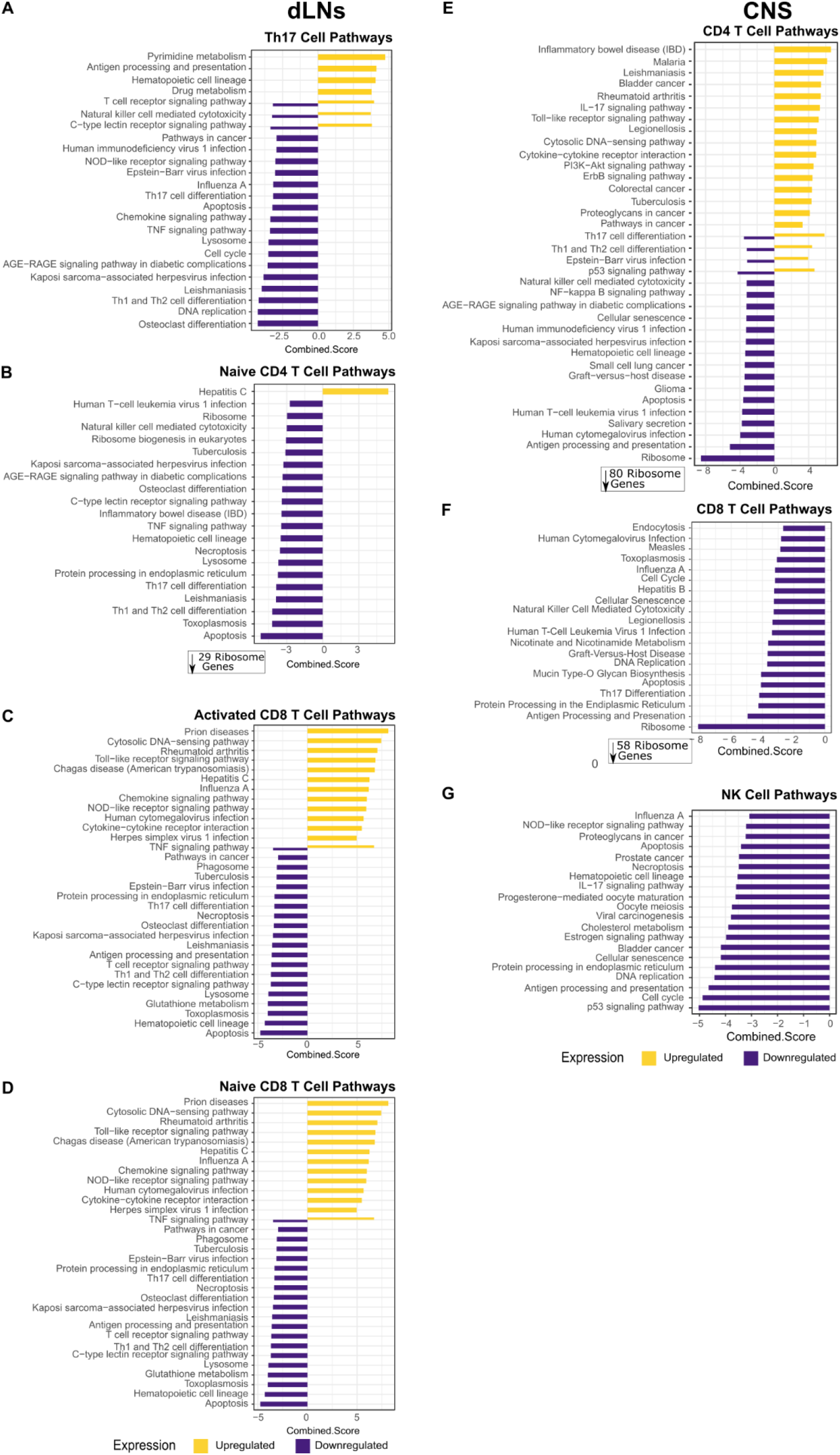
T Cells downregulated Pro-inflammatory and activation pathways in both the dLNs and CNS. Pathways modulated in a) Th17 Cells, b) naïve CD4+ T cells, c) Activated CD8+ T cells and d) naïve CD8+ T cells located in the dLNs at day 2, including ribosome and DNA replication. Pathways modulated in e) CD4+ T cells, f) CD8+ T cells and g) NK cells in the spinal cord at day 7. Genes with padjusted <0.1 were used in pathway analysis. Significant genes were compared to the KEGG pathways and padjusted <0.1 denoted a pathway was up or downregulated. n=2 mice per group in the dLNs and n=1 mouse per group in the CNS.

**Fig. S4.**
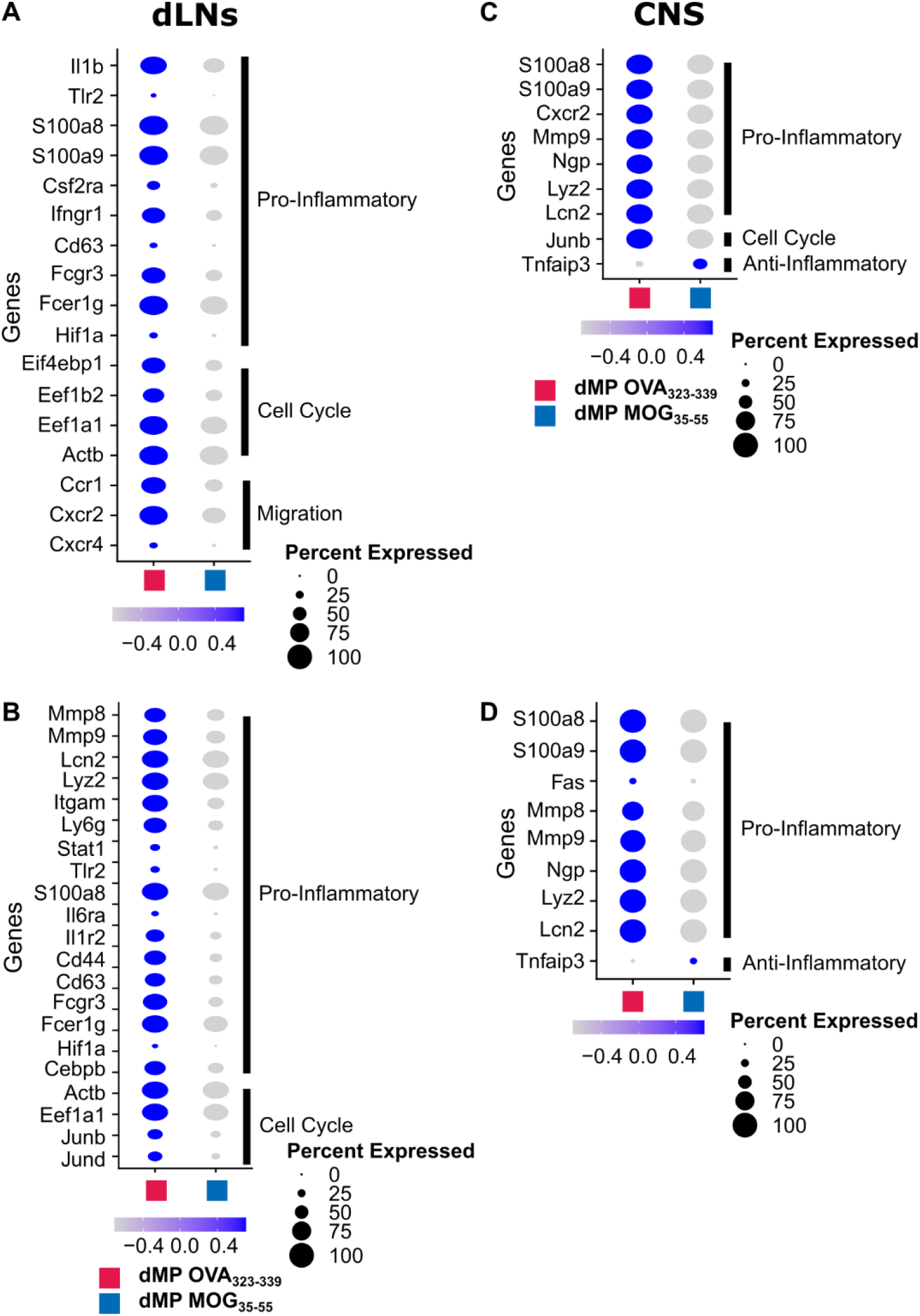
Neutrophils downregulate pro-inflammatory and cell cycle genes in the dLNs and CNS. Dot plots highlighting pro-inflammatory, cell cycle, and migration genes differentially expressed in a) inflammatory neutrophils and b) suppressive neutrophils, specifically genes associated with pro-inflammatory processes in the dLNs at day 2. Dot plots showing downregulation of pro-inflammatory and upregulation of anti-inflammatory genes in the c) inflammatory and d) suppressive neutrophils located in the CNS at day 7. Genes with a p_adjusted_ <0.1 of were considered significant. n=2 mice per group in the dLNs and n=1 mouse per group in the CNS.

**Fig. S5.**
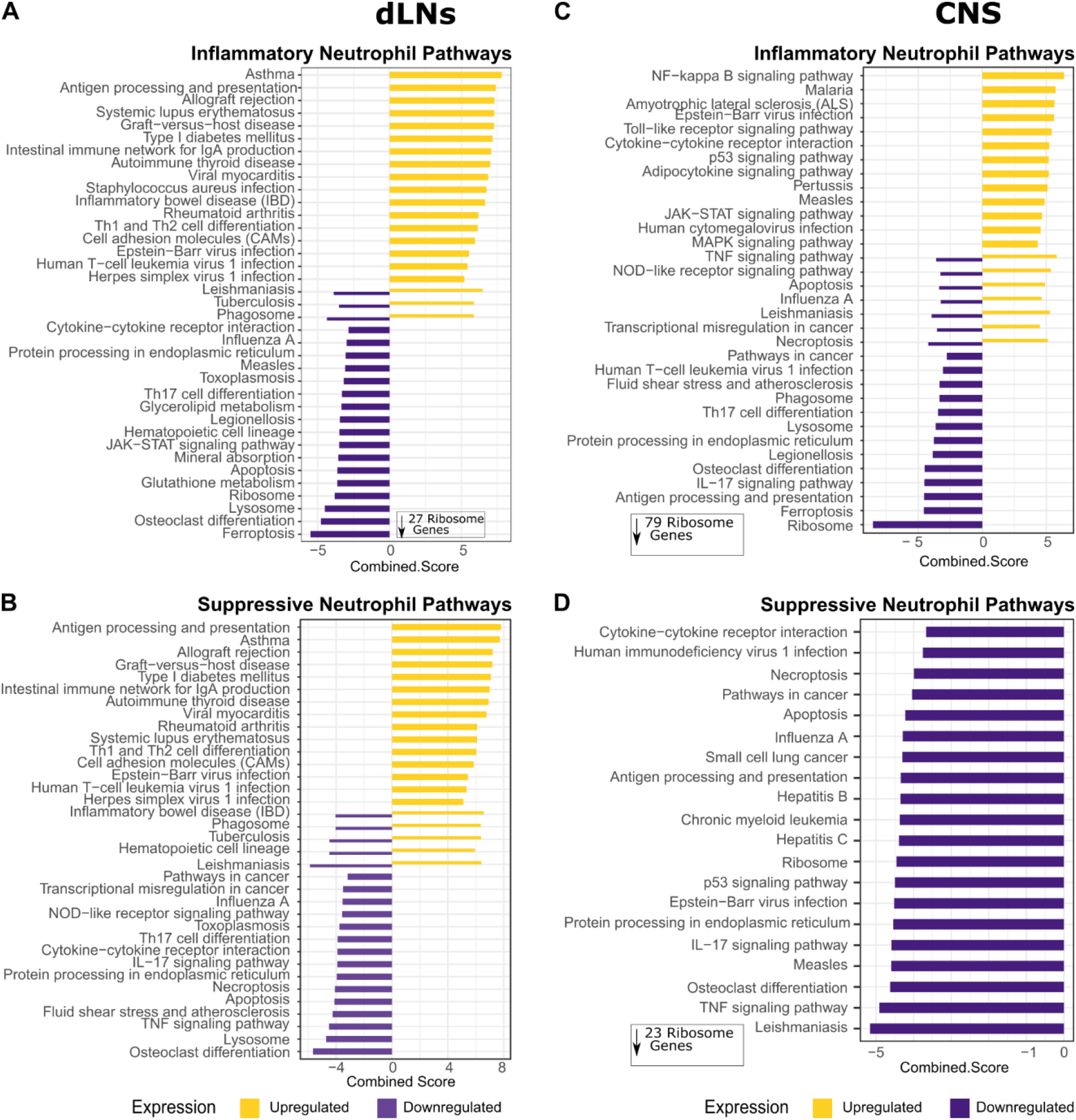
Neutrophils downregulated Pro-inflammatory pathways in both the dLNs and CNS. Pathways modulated in a) inflammatory neutrophils and b) Suppressive neutrophils located in the dLNs at day 2, including ribosome and phagocytosis. Pathways modulated in c) inflammatory neutrophils and d) Suppressive neutrophils located in the CNS at day 7, including ribosome. Genes with p_adjusted_ <0.1 were used in pathway analysis. Significant genes were compared to the KEGG pathways and p_adjusted_ <0.1 denoted a pathway was up or downregulated. n=2 mice per group in the dLNs and n=1 mouse per group in the CNS.

